# ELP1 Gene Augmentation Restores Visual Function in a Mouse Model of Familial Dysautonomia

**DOI:** 10.64898/2026.01.09.698339

**Authors:** Hui-Chen Cheng, Swanand Koli, Kate Lewis, Yasaman Anvarinia, Yating Liu, Morgan Erdoes, Caitlin Elizabeth Keiper, Reynette Estelien, Matthew Chagnon, Luk. H. Vandenberghe, Dadi Gao, Frances Lefcort, Elisabetta Morini, Susan A. Slaugenhaupt, Anil Chekuri

**Affiliations:** Grousbeck Gene Therapy Center, Schepens Eye Research Institute, Massachusetts Eye and Ear Infirmary, Boston, MA, USA; Department of Ophthalmology, Taipei Veterans General Hospital, Taipei, Taiwan; Department of Ophthalmology, College of Medicine, National Yang Ming Chiao Tung University, Taipei, Taiwan; Department of Ophthalmology, Harvard Medical School; Center for Genomic Medicine, Massachusetts General Hospital, Boston, MA, USA; Department of Neurology, Massachusetts General Hospital and Harvard Medical School, Boston, MA, USA; Tikun Therapeutics, 20300 Lubar Way, Brooksville, MD, USA

**Keywords:** Adeno-associated viruses 2, elongator acetyltransferase complex subunit 1, familial dysautonomia, gene therapy, optic neuropathy

## Abstract

Familial dysautonomia (FD) is an autosomal recessive sensory and autonomic neurodevelopmental and degenerative disorder characterized by complex neurological phenotypes. One of its most debilitating features is progressive optic neuropathy, which leads to severe visual impairment in FD patients by the third decade of life. Although several preclinical approaches have shown partial rescue of retinal ganglion cell (RGC) degeneration through increasing Elongator acetyltransferase complex subunit 1 (ELP1) expression in the retina, currently no treatments exist to prevent vision loss in FD. In this study, we performed a comprehensive analysis of visual function in a retina-specific FD mouse model (*Pax6-Cre*_⁺_*;Elp1^loxp/loxp^*) and evaluated a gene supplementation strategy to restore human ELP1 protein levels in the retina. Longitudinal retinal assessments indicated that FD mice exhibit significant retinal nerve fiber layer (RNFL) thinning, as observed in FD patients. FD mice also showed reduced flash visual evoked potentials (VEPs), pattern electroretinography (pERGs), and photopic negative responses (phNRs) amplitudes, along with impaired visual acuity and contrast sensitivity, as assessed using optomotor response assay (OMR). Full-field electroretinography (ffERG) revealed reduced amplitude of dark-adapted a-waves, dark and light-adapted b-waves, indicating combined RGC and bipolar cell dysfunction. Intravitreal delivery of an adeno associated vector (AAV) vector (AAV2.U1a.hELP1) effectively restored physiological ELP1 protein expression, which resulted in a significant rescue of retinal structure and function. Gene supplementation with AAV2.U1a.hELP1 resulted in broad functional and structural improvement compared with untreated FD mice. In summary, our findings provide the first demonstration that ELP1 gene supplementation can effectively rescue RGC function in an FD mouse model and support AAV2.hELP1 at an optimized dose (5.4×10^8^ vg) as a promising therapeutic approach for FD-associated optic neuropathy.

## Introduction

Familial dysautonomia (FD) is a rare, congenital sensory and autonomic neuropathy that affects individuals of Ashkenazi Jewish (AJ) ancestry, with a carrier rate among AJ of 1:32 ^1–6^. FD presents with decreased sensitivity to pain and temperature, cardiovascular instability, and gait ataxia, with an average life span of 40 years^2,4,7^. FD results from a T to C nucleotide change in the 5’ splice site of intron 20 of the Elongator acetyltransferase complex subunit 1 (*ELP1,* formerly known as *IKBKAP*) gene ^8,9^. This mutation leads to tissue-specific skipping of *ELP1* exon 20, resulting in reduced ELP1 protein predominantly in the central and peripheral nervous systems ^8^. ELP1 is a scaffold protein of a highly conserved multiunit Elongator complex that plays a key role in transcription, translation and tRNA modification ^10–14^.

In addition to their peripheral nervous system deficits, FD patients also suffer from progressive visual loss, which severely impacts their quality of life ^3,15–19^. Major ocular defects include decreased visual acuity, loss of color vision, visual field defects and corneal opacities^16,18,19^. FD patients suffer from progressive optic neuropathy beginning as early as five years of age that progresses slowly until approximately 15-20 years of age. Patients then have a rapid visual decline leading to dramatic visual impairment by their second decade of life ^15^. Significant thinning of the retinal nerve fiber layer (RNFL) and ganglion cell-inner plexiform layer (GCIPL) thickness measured by optical coherence tomography (OCT) has been observed in FD patients compared to healthy controls ^18^. The rate of loss was significant before reaching its lowest level between the ages of 24.8 and 26.2 years ^17^. Since FD patients have diminished proprioception, loss of vision severely affects patient mobility, leading to frequent falls, further compromising their quality of life ^3,18^. The slow rate of early progression of optic neuropathy may offer a therapeutic window in which targeted intervention could help prevent eventual visual loss in FD patients.

There are currently no effective treatments for patients with FD. However, several therapeutic strategies are being developed to correct the underlying *ELP1* splicing defect and increase wild-type *ELP1* expression, including splicing modulator compounds (SMCs), antisense oligonucleotides, and gene therapies^20–26^. Some of these approaches have shown partial restoration of *ELP1* levels in the retina and rescue of retinal ganglion cell (RGC) loss with varying degrees of success in preclinical models^20,24,25,27^. Recently, Shultz et al. demonstrated that AAV-mediated supplementation of human *ELP1* reduces RGC loss in a conditional, retina-specific FD knockout mouse model, providing important proof of concept for this therapeutic strategy. However, clinically translatable outcomes, such as retinal nerve fiber layer (RNFL) thickness and visual function, were not evaluated. Here, we performed a comprehensive analysis of retinal structure and visual function in the FD mouse model (*Pax6-Cre*_⁺_;*Elp1^loxp/loxp^*). Furthermore, we demonstrated that AAV-mediated supplementation of human ELP1 (AAV2.U1a.hELP1) effectively restores ELP1 protein expression in the FD retina following intravitreal administration and significantly mitigates optic neuropathy. To our knowledge, this is the first demonstration that visual function in FD can be restored through AAV2-mediated ELP1 gene supplementation

## Methods

### Animal Studies

All mice were maintained in the Schepens Eye Research Institute (SERI) animal facility. All procedures were reviewed and approved by the SERI Institutional Animal Care and Use Committee (SERI-IACUC) of Mass General Brigham (protocol no. 2021N000111) and conducted in accordance with the ARVO Statement for the Use of Animals in Ophthalmic and Vision Research and institutional guidelines. Retina-specific *Elp1* conditional knockout (CKO) mice (*Pax6-Cre*_⁺_*;Elp1^flox/flox*, hereafter referred to as FD mice) were generated as previously described (Ueki *et al.*, 2018)^28^. Briefly, *Elp1 flox/flox* mice (International Knockout Mouse Consortium, Wellcome Sanger Institute, UK) were crossed with *Pax6-Cre* transgenic mice in which Cre recombinase expression is driven by the *Pax6* promoter^28,29^. Both strains were generously provided by Dr. Frances Lefcort (Montana State University). This model enables targeted deletion of *Elp1* in retinal progenitor cells, recapitulating the progressive RGC loss observed in FD while maintaining overall systemic health.

Littermate *Pax6-Cre*_⁻_*;Elp^^flox/flox^* mice were used as controls. Both male and female mice were included in all experiments. Mice were randomly assigned on postnatal day 14 (P14) to either the treatment group or the uninjected control group. For the AAV2.hElp1 treatment studies, a total of seven experimental groups were included: (1) uninjected control; (2) control mice injected with AAV2.hELP1 (5.4 × 10□ vg); (3) control mice injected with AAV2.hELP1 (2.7 × 10□ vg); (4) uninjected FD mice; (5) FD mice injected with AAV2.hELP1 (5.4 × 10□ vg); (6) FD mice injected with AAV2.hELP1 (2.7 × 10□ vg); and (7) FD mice injected with AAV2.eGFP (4.75 × 10□ vg). Visual structure and function were assessed at multiple time points. Spectral-domain optical coherence tomography (SD-OCT) was performed at 1, 3, and 6 months. Electrophysiological assessments including full-field electroretinography (ERG), flash visual evoked potential (VEP), pattern ERG, and photopic negative response (phNR) as well as optomotor response (OMR) testing were conducted at P30, P90 and P180 for all experimental groups. At the conclusion of the study (P180), mice were euthanized, and the retinas were collected for immunohistochemistry (Fig. 1a). A separate cohort of uninjected mice served as a natural history group for both control and FD genotypes, undergoing longitudinal OCT and electrophysiological testing at 1, 3, 6, 12, and 18 months, and OMR testing at 3, 6, 12, and 18 months. The number of mice (n) used in each group is indicated within the bar graphs of the figures. Under standard conditions, mice were maintained in a 12-hour light/dark cycle with *ad libitum* access to food and water. For dark adaptation studies, animals were housed in a dedicated dark chamber, where food and water remained freely available, and were subsequently returned to standard housing. Animal usage was minimized in accordance with both ethical guidelines and experimental design requirements.

**Figure 1:**
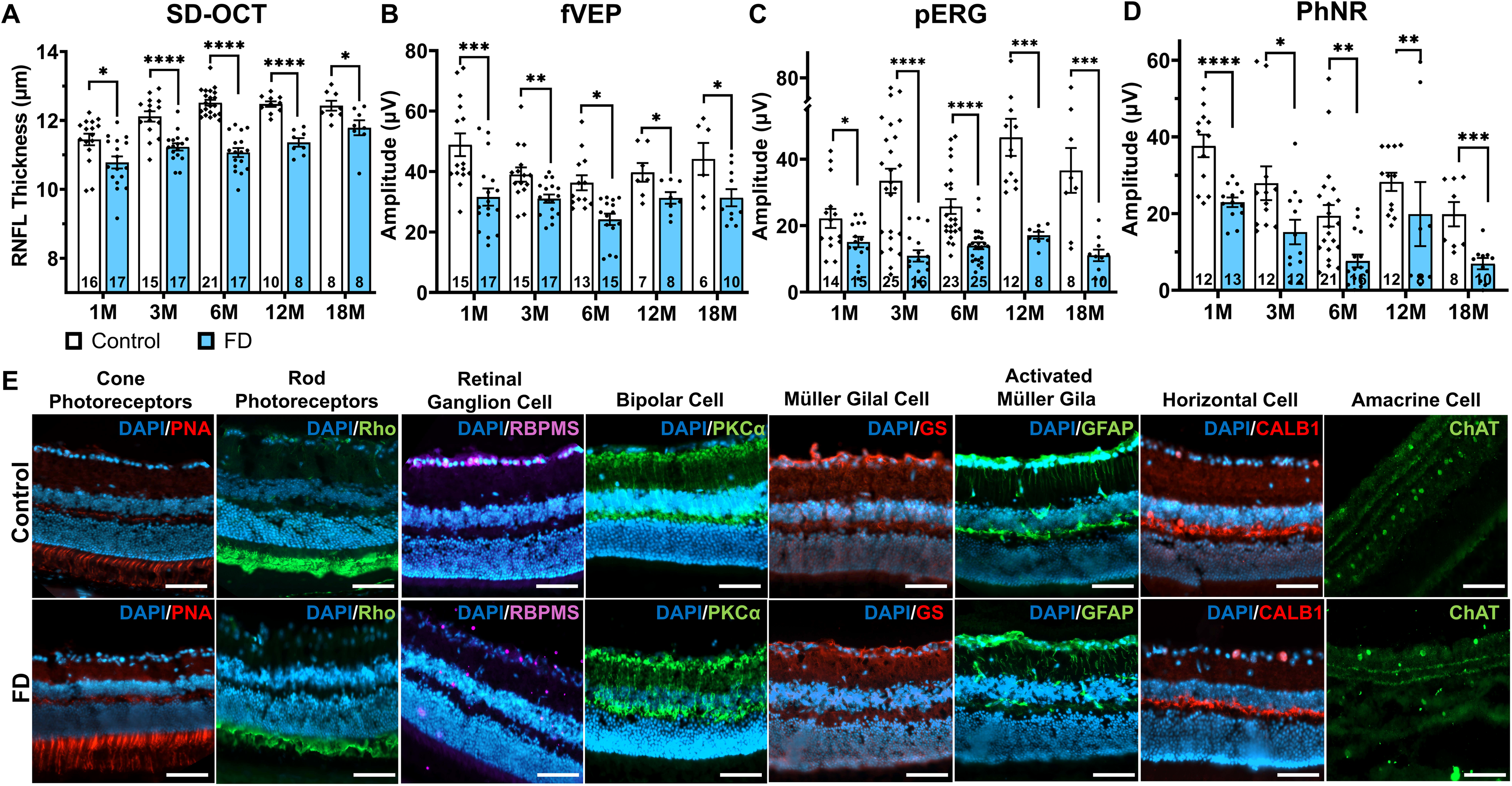
Longitudinal structural, functional, and cellular characterization of retinal degeneration in control and FD mice. (A) Spectral-domain optical coherence tomography (SD-OCT) measurements of retinal nerve fiber layer (RNFL) thickness in 1, 3, 6, 12 and 18 months FD and control mice (B) flash visual-evoked potentials (VEPs), (C) pattern electroretinograms (pERG), and (d) photopic negative responses (phNR) recorded in control and FD mice from 1 to 18 months of age. Data are presented as mean ± SEM and were analyzed using either a t test or Kolmogorov–Smirnov test. Sample sizes for each group are indicated by *n* within each bar. Statistical significance is denoted as *\*p < 0.05, **p < 0.01, ***p < 0.001, ****p < 0.0001*. (e) Representative immunohistochemical analysis of retinal cross sections from control and FD mice. From left to right: peanut agglutinin (PNA) labeling of cone photoreceptors, rhodopsin labeling of rod photoreceptors, RNA-binding protein with multiple splicing (RBPMS) labeling of retinal ganglion cells, protein kinase C alpha (PKCα) labeling of bipolar cells, glutamine synthetase (GS) labeling of Müller glia, glial fibrillary acidic protein (GFAP) labeling of activated Müller glia, calbindin (CALB1) labeling of horizontal cells, and choline acetyltransferase (ChAT) labeling of amacrine cells. Scale bar, 50 µm.

### Intravitreal Injections (IVI)

Postnatal day 14 (P14) mice were anesthetized by intraperitoneal injection of ketamine and xylazine (ketamine 50-75 mg/kg and xylazine 5-7.5 mg/kg mixture). Prior to induction, pupils were dilated using one drop each of 1% tropicamide (Tropicamide Ophthalmic Solution, USP 1%, NDC 24208-585-64; Bausch & Lomb Americas Inc.) and 0.5% proparacaine hydrochloride (Proparacaine Hydrochloride Ophthalmic Solution, USP 0.5%, NDC 24208-730-06; Bausch & Lomb Americas Inc). Under a dissecting microscope, a small puncture was created in the sclera using a 30-gauge needle. A 33-gauge blunt-tip needle attached to a 10-µL Hamilton syringe was then inserted through the puncture site at a perpendicular angle to the eye surface. The needle was advanced through the sclera and choroid into the vitreous chamber, and 1 µL of AAV solution was slowly injected into the vitreous cavity. Following injection, the needle was held in place for 30–40 seconds to minimize reflux, then slowly withdrawn, and the conjunctiva gently repositioned. A topical ophthalmic lubricant (Genteal, Alcon Laboratories) was applied to prevent corneal drying during recovery. Each mouse was considered an independent experimental unit. Intravitreal injections were performed using AAV2.U1a.hELP1, carrying either 5.4 × 10□ or 2.7 × 10□ vector genomes (vg) of human *ELP1* (hELP1), driven by the murine small nuclear RNA promoter (*U1a*) and expressing the full-length human *ELP1* sequence (Origene RC2076868)^24^. For control-treated FD eyes, AAV2.eGFP (generated at Gene Therapy Vector Core, SERI) at 4.75 × 10□ vg was injected under identical conditions. The AAV2.U1a.hELP1 was also generously gifted by Dr. Frances Lefcort (Montana State University).

### Spectral Domain–Optical Coherence Tomography (OCT)

Mice were anesthetized with ketamine and xylazine prior to imaging. Retinal morphology was assessed using a Spectral Domain–Optical Coherence Tomography (SD-OCT) system (Envisu R2210 UHR, Leica Microsystems, USA) equipped with a 50° field of view. The scan was centered on the optic nerve head and acquired using a rectangular volume scan covering 1.4 mm × 1.4 mm with 1000 A-scans × 100 B-scans × 3 repeats. OCT images from each eye were averaged and analyzed using InVivoVue (version 2.4.35) and InVivoVue Diver (version 3.4.4) software (Bioptigen, USA). Retinal thickness was measured and segmented using the automated segmentation algorithm, followed by manual verification when necessary. The segmented retinal layers were grouped into three regions for analysis: the retinal nerve fiber layer (RNFL), the middle retinal layers comprising the inner plexiform layer (IPL), inner nuclear layer (INL), and outer plexiform layer (OPL), and the outer retinal layers including the outer nuclear layer (ONL), inner segments (IS), and outer segments (OS). All quantitative data are expressed as mean ± SEM.

### Full-field electroretinography (ffERG)

For all electrophysiological recordings, mice were dark-adapted overnight. On the day of examination, animals were anesthetized with ketamine and xylazine, and body temperature was maintained at 37°C using a feedback-controlled heating pad throughout the procedure. Pupils were dilated with 1% tropicamide and 2.5% phenylephrine before recording. Measurements were obtained from both eyes, and each eye was considered an independent experimental unit unless otherwise specified. ERG recordings were obtained using corneal contact gold wire electrodes with a reference electrode placed subcutaneously between the ears and a ground electrode at the base of the tail. Signals were amplified (gain ∼1,000×), bandpass filtered (0.1–500 Hz), and digitized using an acquisition system (Espion V6.68.2, Diagnosys, USA). For scotopic ERG, a series of increasing flash intensities (from 0.000249 to 24.1 cd·s/m²) were delivered using a Ganzfeld dome (Color Dome LabCradle, Diagnosys, USA) under dark-adapted conditions to elicit rod-dominated responses. A-wave and b-wave amplitudes were measured as the first negative and subsequent positive peaks, respectively. After light adaptation to a white background light (30 cd/m²) for 7 minutes, cone-driven responses were recorded using 2 ms white light flashes (0.1-25.6 cd·s/m²). For phNR, white flash with intensity of 10 cd.s/m^2^ was used to elicit a robust negative-going wave following the b-wave. The phNR amplitude was measured from the baseline to the trough following the b-wave. At least 75 responses were averaged per condition to improve signal-to-noise ratio.

### Pattern electroretinography (pERG)

Pattern ERG signals were recorded using a commercially available system (Celeris, Diagnosys, USA) with a visual pattern stimulus consisting of alternating black and white horizontal gratings (spatial frequency: 0.0589 cycles/degree; contrast: 100%) displayed on a pattern electrode positioned on the animal’s eye. The stimulus was presented in a pattern-reversal mode at a frequency of 2 Hz without a mean luminance change. The pattern electrode on each eye served as the active electrode. We used a monocular protocol, which stimulates one eye at a time and uses the pattern electrode on the opposite eye as the reference and ground. Signals were amplified (×10,000), bandpass filtered (1–50 Hz), and averaged over 400 sweeps to enhance signal-to-noise ratio, using an acquisition system (Espion software V6.68.2, Diagnosys, USA). PERG responses were analyzed by measuring the amplitude of the P1–N2 component, defined as the peak-to-trough difference between the first positive (P1) and subsequent negative (N2) waves.

### Flash visual-evoked potential (fVEP)

For flash VEP, a subdermal stainless-steel electrode was placed at the midline over the location of the visual cortex as the active electrode, with reference and ground electrodes placed at the snout and the base of the tail. Visual flash stimuli consisted of brief white light flashes (duration: 1–2 ms; intensity: 3 cd·s/m²) delivered from light-guide electrodes under light-adapted conditions. Each eye was stimulated separately by the light-guide electrode on each eye. Electrophysiological signals were amplified (gain ∼10,000×), bandpass filtered (0.125-300 Hz), digitized, and averaged across 375 trials using a data acquisition system (Espion V6.68.2, Diagnosys, USA). Flash VEP responses were analyzed by measuring the amplitude of the P2–N1 component, defined as the peak-to-trough difference between the first negative deflection (N1) and the second positive (P2) peaks. All the measurements were done using Espion V6.68.2 (Diagnosys, USA).

### Optomotor response (OMR)

The optomotor response (OMR) was evaluated using a virtual optokinetic drum (OptoDrum, Striatech GmbH, Germany) to assess spatial visual acuity (VA) and contrast sensitivity (CS) in mice. Animals were placed individually on an elevated central platform surrounded by a rotating virtual cylinder displaying vertical sine wave gratings. Stimuli were presented in alternating clockwise and counterclockwise directions to preferentially stimulate each eye. Testing was conducted under uniform ambient illumination (∼60 lux) during the light phase. For visual acuity (VA) assessment, the grating contrast was fixed maximum contrast (99.72%), and the spatial frequency was progressively increased in 0.02 cycles per degree (cpd) increments until no consistent tracking movement was observed. The highest spatial frequency eliciting a reliable head-tracking response was recorded as the VA threshold.

For contrast sensitivity testing, the spatial frequency was fixed at 0.061, 0.133, 0.208, 0.267, 0.389 cpd, respectively. The contrast decreased stepwise from 99.72 (maximum) to 0.86% until the OMR was no longer detectable. The inverse of the minimum contrast threshold was defined as the contrast sensitivity. OMR behavior was automatically detected and verified by an observer blinded to experimental groups using the OptoDrum software suite.

### Retinal Whole-Mount Staining and RGC Counting

To quantify the RGC number and transgene expression, we performed immunofluorescence staining and quantitative analysis on retinal whole mounts. Our protocol was adapted from previously established methods (Ueki et al.) with slight modifications to enable multiplex labeling^28^. Mice were euthanized by CO₂ asphyxiation, and eyes were promptly enucleated. Eyes were fixed in 4% paraformaldehyde (PFA) (Electron Microscopy Sciences #15714-S) for 16 hours at 4°C. Following fixation, retinas were carefully dissected free of the cornea, lens, vitreous, and sclera. Four radial relieving cuts were made to flatten each retina for whole-mount processing. To minimize non-specific antibody binding, retinas were incubated overnight at 4°C in a blocking solution consisting of 1X Goat blocking serum (1X GBS) with 0.5% Triton X-100. Primary antibody incubations were performed for 48-72 hours at 4°C. To quantify all RGCs, a rabbit anti-RBPMS antibody (PhosphoSolutions #1830-RBPMS; 1:500) was used. AAV-mediated GFP transgene expression was visualized using a chicken anti-GFP antibody (1:1000) to detect the reporter and a rabbit anti-hIKAP antibody (Thermo Fisher# PA5-111296; 1:500) to detect the overexpressed human ELP1 protein. Select retinas were co-stained with a mouse anti-Brn3a antibody ((Santa Cruz Biotechnology# sc-8429 AF488;1:200) for complementary RGC labeling. Following three 5-minute washes in 1X DPBS, retinas were incubated for 90 minutes at room temperature with appropriate secondary antibodies conjugated to Alexa Fluor dyes (Donkey anti-rabbit 568, Donkey anti-chicken 488, Goat anti-mouse 488; all used at 1:1000 in 1X DBS). Finally, retinas were mounted with retinal ganglion cell layer up on slides using VECTASHIELD Antifade Mounting Medium (Vector Laboratories #H-2000-10). Supplementary table 1 enlists all the antibodies used in the experiment. High-resolution imaging of entire retinal whole mounts was performed using a Zeiss Axio Imager M2 Upright Fluorescence epifluorescence microscope integrated with a Tissue Gnostics Tissue FAXS SL Q system available at Ragon Institute, Massachusetts General Hospital. The high-throughput scanning confocal system was used to acquire tiled, optical-section edges (z-stacks) at 20X magnification. For each tile, a z-stack with a step interval of 4 µm top and below was captured to ensure all fluorescent cells within the volume of the tissue were recorded. Automated stage and autofocus functions of the system were used to generate composite images (whole-retinal scans) encompassing approximately 12-15 mm² of each retina, which were then flattened into a maximum intensity projection for analysis.

Quantification of RBPMS-positive RGCs was performed using a customized automated analysis pipeline developed in CellProfiler to ensure objective and reproducible measurements. The workflow included correction for uneven illumination across the composite images, followed by automatic detection of cell-sized objects based on typical RGC diameter and fluorescence intensity in the RBPMS (Alexa Fluor 568) channel. Minimum intensity thresholds were applied to distinguish true RBPMS-positive cells from background autofluorescence or nonspecific signal. Cell counts were obtained from four defined 1 mm² regions of interest positioned 1 mm from the optic nerve head in the superior, inferior, nasal, and temporal quadrants. If a primary region was damaged (e.g., by a tissue fold or tear), an adjacent undamaged area of equal size within the same quadrant was analyzed. All image acquisition and CellProfiler analyses were conducted by an investigator blinded to treatment groups. Statistical analyses were performed using GraphPad Prism (v10.6.1).

### Immunohistochemical analysis (IHC)

To evaluate potential cellular and structural changes following AAV2-mediated hELP1 overexpression, we performed immunofluorescence analysis on retinal sections using a panel of well-characterized cell-type-specific markers. Eyes from both AAV2-hELP1 injected mice and uninjected control mice were enucleated immediately following CO_2_ asphyxiation. They were fixed by immersion in 4% paraformaldehyde (PFA) in 0.1 M phosphate buffer saline w/o calcium & magnesium (1x DPBS, pH 7.4) for 2-4 hours at 4°C. After fixation, eyes were cryoprotected by sequential immersion in 15% and 30% sucrose solutions in 1x DPBS overnight at 4°C. Eyes were then embedded in optimal cutting temperature (OCT) compound (Scigen #4586), and 12-μm thick sagittal sections, centered on the optic nerve head, were cut using a cryostat (Leica #CM1950). Sections were mounted on Superfrost Plus slides (Fischerbrand # 1255015) and stored at −80°C until staining.

All staining procedures were performed at room temperature unless otherwise specified. Retinal sections were first bordered with a hydrophobic barrier pen. Sections were rehydrated and permeabilized in 0.3% Triton X-100 in 1x DPBS for 15 minutes, followed by blocking for 1 hour in a solution containing 5% normal goat serum and 1% bovine serum albumin (BSA) in PBS. For optimal epitope detection of specific markers, antigen retrieval was performed prior to immunostaining. Sections designated for staining with anti-RBPMS, anti-PKCα, or anti-GFAP were immersed in pre-heated 10 mM sodium citrate buffer (pH 6.0) and subjected to heat-induced epitope retrieval using a standard steamer method for 20 minutes, followed by a 30-minute cooling period at room temperature. All sections were then permeabilized with 0.3% Triton X-100 in PBS for 15 minutes and blocked for 1 hour in a solution containing 5% normal goat serum and 1% bovine serum albumin in PBS.

Primary antibodies, diluted in blocking solution, were applied overnight at 4°C. We used the following panel to label distinct retinal cell populations: Photoreceptors: Peanut Agglutinin (Invitrogen Lectin PNA, Alexa Fluor™ 568 Conjugate, L32458, 1:200) to label cone sheaths, and mouse anti-Rhodopsin (Abcam, AB5417, 1:500) to label rod outer segments. Horizontal Cells: Rabbit X-Calbindin D-28K (Millipore, AB1778, 1:1000). Bipolar Cells: Mouse anti-PKCα (BD Transduction Laboratories, #610108, 1:200). Amacrine Cells: Goat anti-Choline Acetyltransferase (ChAT; Millipore, AB144P, 1:100). Retinal Ganglion Cells: Rabbit anti-RBPMS (PhosphoSolutions, 1830-RBPMS, 1:500).Müller Glia: Mouse anti-Glutamine Synthetase (GS; Millipore Sigma, G2781-100UL, 1:500) to label all Müller cells, and rabbit anti-Glial Fibrillary Acidic Protein (Davis/NIH Neuro Mab facility, Clone N206A/8,1:500) to label activated Müller glia and astrocytes. The following day, sections were washed three times in 1X DPBS and incubated for 2 hours with species-appropriate secondary antibodies (all at 1:1000 dilution in blocking buffer). The secondary antibodies used (detailed in Supplementary Table S2) were: goat anti-rabbit IgG-Alexa Fluor 568 (Thermo Fisher, #A27040) for RBPMS, Calbindin, GFAP, and hIKAP; goat anti-mouse IgG-Alexa Fluor 488 (Thermo Fisher, #A-11029) for Rhodopsin, PKCα, and GS; donkey anti-goat IgG-Alexa Fluor 488 (Thermo Fisher, #A-11055) for ChAT; and goat anti-chicken IgY-Alexa Fluor 488 (Thermo Fisher, #A-11039) for GFP. Following secondary incubation, sections were washed, and coverslips were applied using VECTASHIELD Antifade Mounting Medium (Vector Laboratories #H-2000-10). Images were captured using a Zeiss Axio Imager M2 Upright Fluorescence Motorized Microscope. For consistent analysis, images from the central retina (adjacent to the optic nerve head) were acquired using a 20x objective with identical laser power, gain, and pinhole settings across all samples within an experiment. All images were processed using ZEISS ZEN 3.5 (blue edition) software.

### Statistical analysis

The Shapiro-Wilk test was used to assess whether the data followed a normal distribution. For normally distributed continuous variables, Student’s t-test was applied; otherwise, the Kolmogorov-Smirnov test was used. To compare multiple experimental groups, analysis of variance (ANOVA) with post hoc multiple comparisons was performed for normally distributed data. All statistical analyses were conducted using GraphPad Prism version 10.1.1 for macOS (GraphPad Software). A two-sided p-value < 0.05 was considered statistically significant. The complete set of original *p*-values for all experiments is included in Supplementary Table S2.

## Results

### Comprehensive characterization of retinal degeneration in a retina-specific FD mouse model

Previous work by Ueki et al. characterized the retinal histopathology of the FD mouse model^28^. However, a comprehensive evaluation of the structural and functional defects in this model has not been performed. Such analysis is essential not only for understanding the progression and mechanisms of retinal pathology in FD but also for establishing clinically relevant endpoints for future therapeutic studies. To define the natural history of retinal degeneration in vivo in a retina-specific FD mouse model, we performed comprehensive longitudinal structural, functional, and cellular assessments at 1, 3, 6, 12, and 18 months of age. SD-OCT imaging revealed a reproducible and progressive pattern of retinal pathology in FD mice, characterized by significant thinning of the retinal nerve fiber layer (RNFL) across all time points (*p<0.05-0.0001*) (*Figures 1A*). These structural abnormalities emerged early and worsened with age, consistent with progressive RGC loss observed in FD patients. A comprehensive visual function analysis of the FD retina corroborated the structural abnormalities. FD mice exhibited significantly reduced flash visual evoked potentials (VEPs), pattern electroretinogram (pERG) responses, and photopic negative response (phNR) amplitudes at all examined ages ( *p<0.05)*, reflecting impaired RGC signaling and dysfunction of downstream visual pathway (*Figures 1B-D*) To determine whether the structural and functional abnormalities observed in the FD retina corresponded with underlying cellular changes, we performed a comprehensive immunohistochemical analysis of all major retinal cell classes. Staining of retinal cross-sections confirmed the physiological deficits measured by OCT and electrophysiology, revealing a pronounced and selective loss of RGCs in FD mice. In contrast, other retinal layers, including photoreceptors, bipolar cells, amacrine cells, and Müller glia showed largely preserved morphology and cell number (Figure. 1E). These findings indicate that Elp1 deficiency leads to a highly selective pattern of RGC degeneration, while sparing the broader retinal architecture. Furthermore, behavioral testing using the optomotor response (OMR) assay revealed significantly reduced visual acuity and contrast sensitivity in FD mice across multiple spatial frequencies and ages (*p<0.05*), consistent with the observed structural and electrophysiological deficits and confirming impaired visually driven behavior (*Figures 2B–G*). Given the structural degeneration observed in FD retinas in our OCT analysis, we next examined whether *ELP1* deficiency leads to measurable changes in photoreceptor and inner retinal function. Full-field ERG recordings provided additional insights into outer and inner retinal integrity. In the natural history cohort, control mice exhibited lower dark-adapted *a*-wave amplitudes under high-intensity stimuli compared with FD mice at all time points except 12 months (*Figure 2Ab*) *(p<0.05)*. No significant differences were observed in light-adapted *a*-wave amplitudes, except for slightly reduced amplitudes in control mice under higher stimulus intensities at 1 and 18 months. In contrast, FD mice consistently showed lower *b*-wave amplitudes under both dark- and light-adapted conditions compared with FD mice across all examined ages *(p<0.05)* (*Figure 2A*). Taken together, the results demonstrate significant impairment of photoreceptor and inner retinal function in FD mice relative to controls.

**Figure 2:**
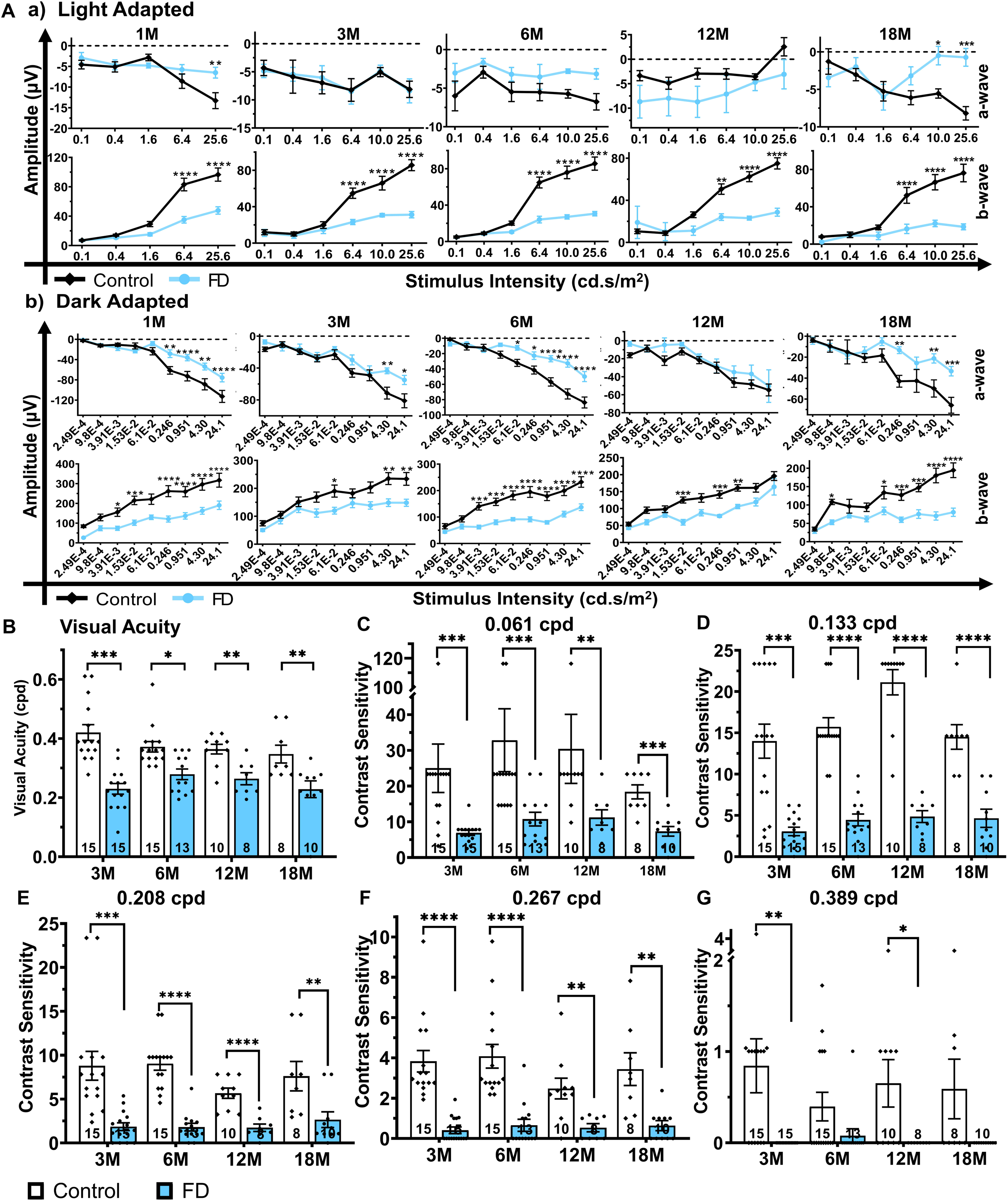
Age-dependent changes in ERG responses and visual behavior in FD mice. (A) (a) Light-adapted and (b) dark-adapted full-field electroretinogram (ERG) responses recorded in control and FD mice from 1 to 18 months of age, revealing progressive functional deficits in FD mice. Data are presented as mean ± SEM and were analyzed by two-way ANOVA; *p* < 0.05, p < 0.01, *p* < 0.001, p < 0.0001. (B) Visual acuity measured by optomotor response (OMR) in control and FD mice from 3 to 18 months of age. (C–G) Contrast sensitivity assessed at spatial frequencies of 0.061, 0.133, 0.267, 0.389, and 0.208 cycles per degree (cpd), respectively, demonstrating age-dependent impairments in visual performance in FD mice. Data are presented as mean ± SEM and were analyzed using either a t test or a Kolmogorov–Smirnov test. Sample sizes for each group are indicated by *n* within each bar; *\*p < 0.05, **p < 0.01, ***p < 0.001, ****p < 0.0001*.

### AAV2.hELP1 treatment modulates RNFL thickness in FD mice

To determine whether intravitreal delivery of AAV2.hELP1 can prevent retinal degeneration we performed intravitreal injections in FD and control mice at postnatal day 14 (P14), and retinal structure and visual function were assessed at 3 (P90), and 6 months (P180) of age *(Figure 3 A,B*). We first investigated whether intravitreal delivery of AAV2.hELP1 could prevent or reverse RNFL thinning in FD mice. FD mice that received an intravitreal injection of AAV2.hELP1 (2.7 × 10□ vg) exhibited a significantly lower RNFL thickness compared to uninjected FD mice at 1 month (*p < 0.001; Figure S1A*), reflecting an acute response to intravitreal AAV delivery or early-stage disease progression that is not yet fully mitigated by ELP1 supplementation. At 3 months, no significant differences in RNFL thickness were detected among the AAV2.hELP1 treated FD mice compared to the untreated control (*Figure 3Ca*), suggesting that hELP1-mediated rescue has a delayed therapeutic onset and is modulated by the underlying stage of disease progression. By 6 months, FD mice treated with AAV2.hELP1 (5.4 × 10□ vg) or AAV2.hELP1 (2.7 × 10□ vg) displayed a significant increase in RNFL thickness compared with uninjected FD mice (*p < 0.05 and p < 0.0001, respectively; Figure 3Cb*). Together, these findings indicate that AAV2.hELP1 treatment dynamically modulates RNFL thickness in FD mice in a dose and age-dependent manner.

**Figure 3:**
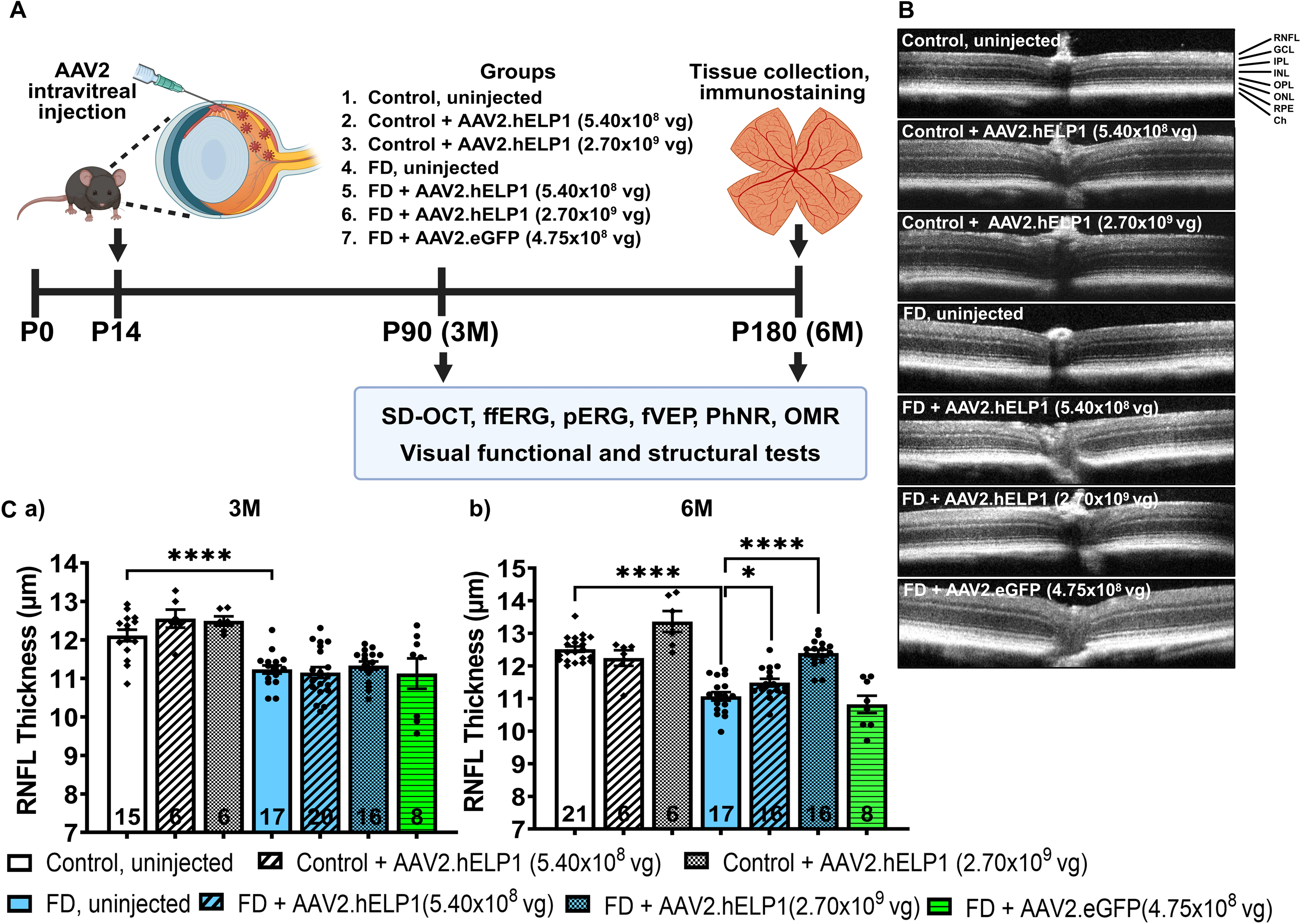
Optical coherence tomography analysis following AAV-mediated gene therapy in FD mice. (a) Schematic of the experimental design. Control and FD mice received bilateral intravitreal injections at postnatal day 14 (P14) of AAV2.hELP1, AAV2.eGFP, or no injection. Retinal structural and visual functional assessments were performed at P90 and P180. At the study endpoint (P180), mice were euthanized and retinas were collected for immunohistochemical analysis. (b) Representative spectral-domain OCT (SD-OCT) images from control non-injected mice, FD non-injected mice, FD mice treated with AAV2.hELP1 (5.4 × 10□ vg), FD mice treated with AAV2.hELP1 (2.7 × 10□ vg), and FD mice treated with AAV2.eGFP (4.75 × 10□ vg). (c) Quantification of retinal nerve fiber layer (RNFL) thickness measured by OCT at 3 and 6 months of age across treatment groups, indicating partial structural preservation following AAV2.hELP1 treatment. Data are presented as mean ± SEM and were analyzed using either a t test or Kolmogorov–Smirnov test. Sample sizes for each group are indicated by *n* within each bar; *\*p < 0.05, **p < 0.01, ***p < 0.001, ****p < 0.0001.* The control group and FD group shown here are the same as that presented in Figure 1 to allow for direct comparison

### Improvement of RGC-related electrophysiological responses following AAV2.hELP1 treatment

To determine whether AAV2.hELP1 gene supplementation restores RGC-mediated visual pathway function, we next evaluated electrophysiological responses in treated and untreated FD mice. Following intravitreal delivery of AAV2.hELP1, we observed significant improvements in electrophysiological responses in treated FD mice. At 3 months and 6 months, FD mice receiving intravitreal AAV2.hELP1 at 5.4 × 10□ vg exhibited significantly higher flash VEP (fVEP) amplitudes compared with uninjected FD mice (*p<0.05; Figure 4A*), suggesting that restoring ELP1 expression improves the electrophysiological responses through the whole visual pathway. At 3 months, this treatment group also showed significantly increased pERG and phNR amplitudes *(p<0.05); Figures 4Bb and 4Cb)*, indicating early recovery of RGC-specific function. By 6 months, both the 5.4 × 10□ vg and 2.7 × 10□ vg AAV2.hELP1 treatment groups demonstrated significantly higher pERG amplitudes than uninjected FD mice *(p < 0.001 and p < 0.0001, respectively; Figures 4Bc and 4Cc)*. Notably, only the lower-dose group (5.4 × 10□ vg) showed a significant enhancement in phNR amplitude at this time point *(p < 0.001; Figure. 4Cc)*, further highlighting a dose-dependent and durable therapeutic effect on RGC function. Overall, these results demonstrate that hELP1 gene supplementation can restore key RGC related visual responses in a dose- and time-dependent manner in FD mice.

**Figure 4:**
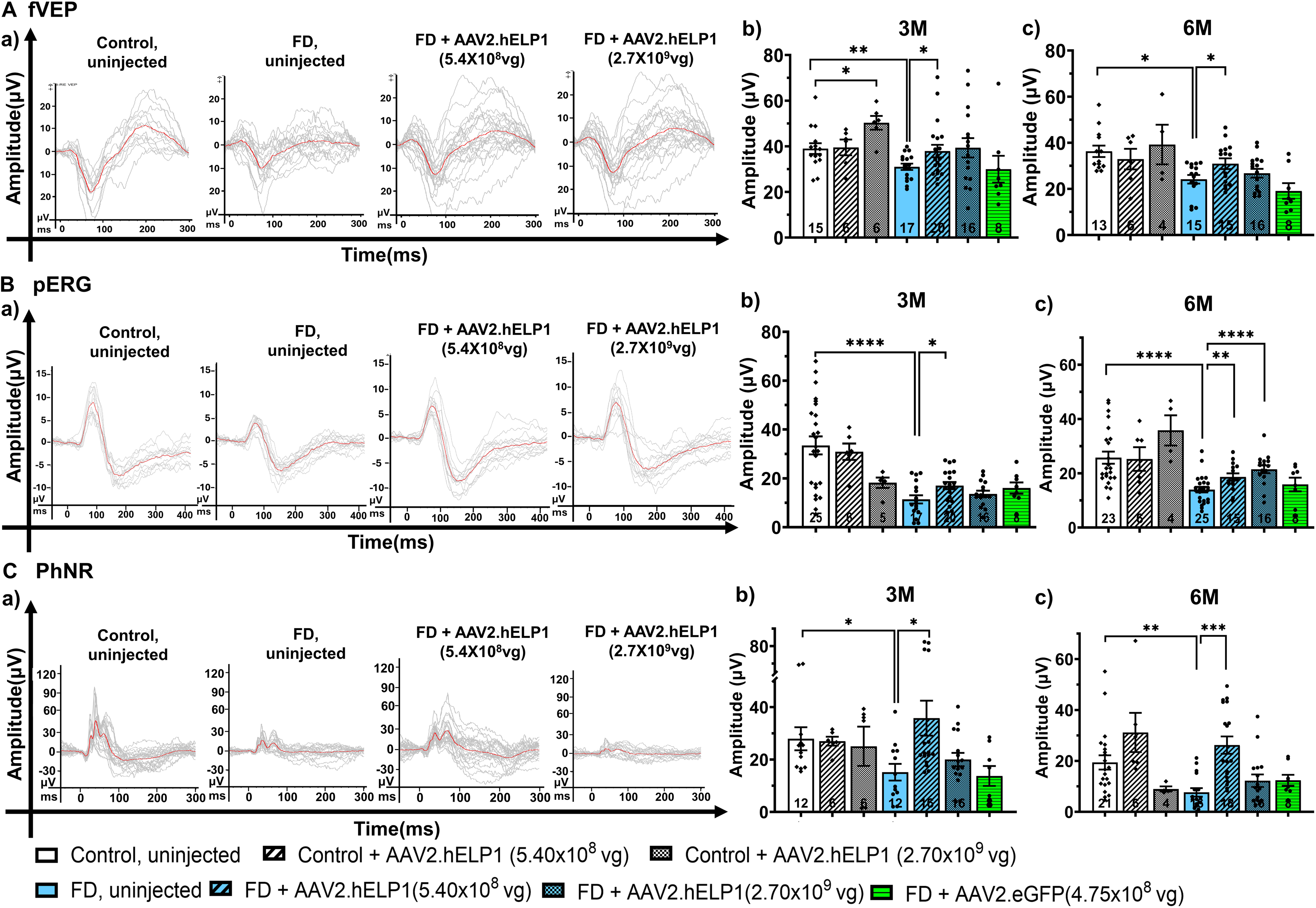
Visual pathway and retinal ganglion cell functional outcomes following AAV-mediated gene therapy in FD mice. (A) (a) Representative waveforms of flash visual-evoked potentials (VEPs) recorded from control mice, uninjected FD mice, and FD mice treated with AAV2.hELP1 at doses of 5.4 × 10□ vg or 2.7 × 10□ vg. (b,c) Quantification of flash VEP P2–N1 amplitudes across all groups at 3 and 6 months, respectively. (B) (a) Representative pattern electroretinogram (pERG) waveforms from all experimental groups. (b,c) Quantification of pattern ERG P1–N2 amplitudes at 3 and 6 months, respectively. (C) (a) Representative photopic negative response (phNR) waveforms recorded from each experimental group. (b,c) Quantification of phNR amplitudes at 3 and 6 months, demonstrating partial functional improvement of inner retinal and visual pathway responses following AAV2.hELP1 treatment. Data are presented as mean ± SEM and were analyzed using either a t test or Kolmogorov–Smirnov test. Sample sizes for each group are indicated by *n* within each bar; *\*p < 0.05, **p < 0.01, ***p < 0.001, ****p < 0.*0001. The control group and FD group shown here is the same as that presented in Figure 1 to allow for direct comparison

### AAV2.hELP1 treatment partially restores inner retinal signaling in FD mice

To determine whether AAV2.hELP1 treatment could improve inner retinal function, we next assessed full-field ERG responses in treated and untreated FD mice. In a typical full field ERG (ffERG), the a-wave represents the initial negative deflection and primarily reflects photoreceptor hyperpolarization, whereas the b-wave is the subsequent positive deflection generated mainly by ON bipolar cells with contributions from Müller glia, serving as a functional readout of inner retinal activity^30–32^. FD mice receiving intravitreal injection (IVI) of AAV2.hELP1 (5.4 × 10□ vg) exhibited better dark-adapted *a*-wave amplitudes but greater dark and light-adapted *b*-wave amplitudes compared with uninjected FD mice at 3 and 6 months (*p<0.05; Figure 5B*). Conversely, FD mice treated with AAV2.hELP1 (2.7 × 10□ vg) showed worse dark-adapted *a*-wave amplitudes and lower *b*-wave amplitudes under both dark- and light-adapted conditions at 6 months (*Figure. 5B*). Similarly, in control mice, intravitreal administration of AAV2.hELP1 (2.7 × 10□ vg) resulted in significantly worse dark-adapted *a*-wave amplitudes and lower *b*-wave amplitudes under both conditions compared with uninjected controls at 6 months (*Figure. 5B*). Together, these full-field ERG findings indicate that *Elp1* deficiency in the retina leads to impaired inner retinal activity in FD mice. Importantly, AAV2-mediated *ELP1* supplementation partially restored inner retinal function, suggesting improved signal transmission from photoreceptors to downstream retinal neurons.

**Figure 5:**
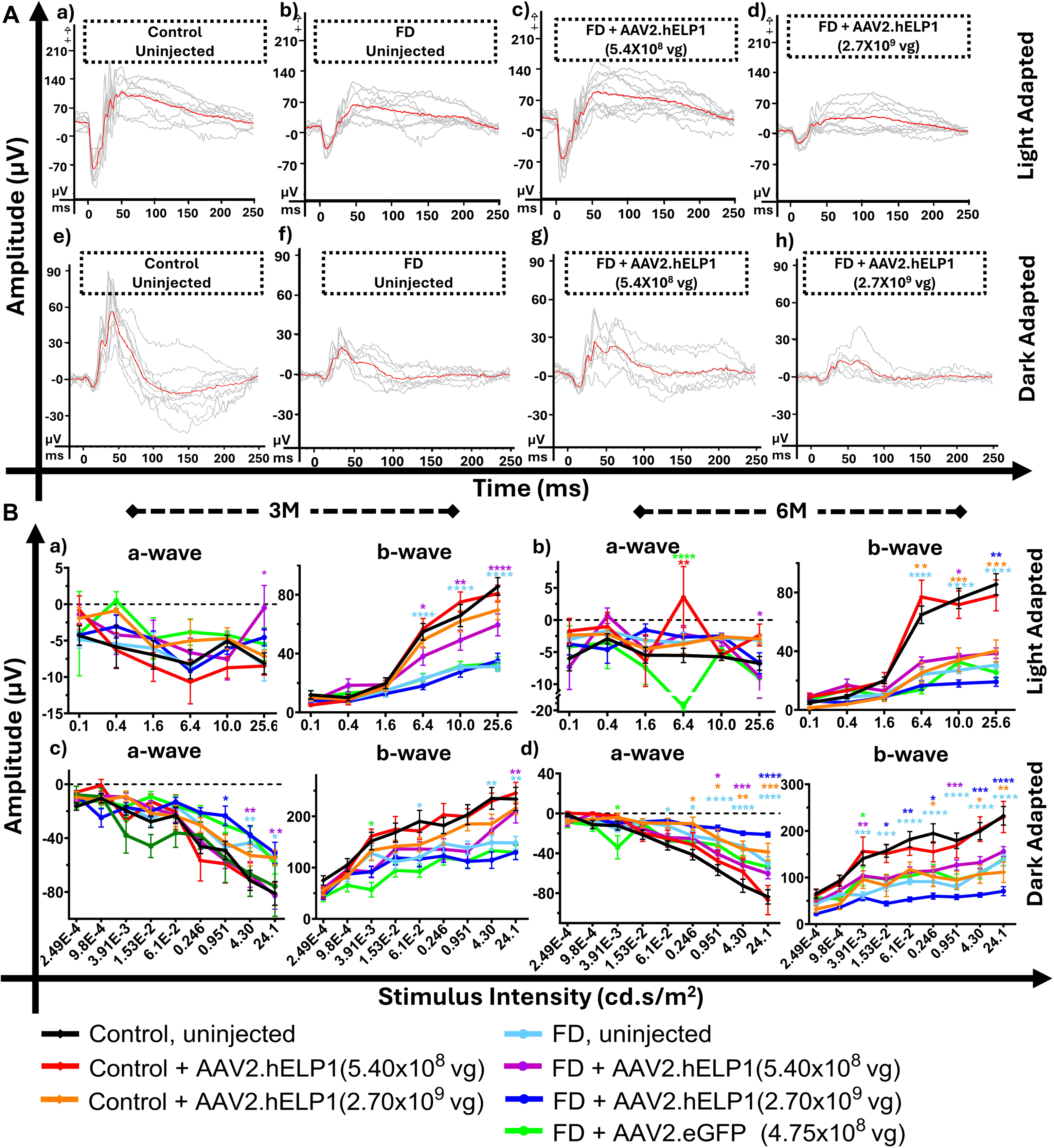
Full-field electroretinographic assessment following AAV-mediated gene therapy in FD mice. A. Representative (a–d) light-adapted and (e–h) dark-adapted ffERG recordings across experimental groups. Light-adapted ffERG responses were recorded at 25.6 cd·s/m² and dark-adapted ffERG responses at 24.1 cd·s/m². Traces represent group-averaged responses, with the mean waveform for each group shown in red. B. (a,b) Quantification of light-adapted a-wave and b-wave amplitudes and (c,d) dark-adapted a-wave and b-wave amplitudes measured at 3 and 6 months of age across all groups, consistent with partial restoration of photoreceptor-driven responses following AAV2.hELP1 treatment. Data are presented as mean ± SEM and were analyzed by two-way ANOVA. Sample sizes for each group are indicated by *n* within each bar; *\*p < 0.05, **p < 0.01, ***p < 0.001, ****p < 0.*0001. The color of the significance markers denotes the comparison group as follows: red, control uninjected vs. control + AAV2.hELP1 (5.4 × 10□ vg); orange, control uninjected vs. control + AAV2.hELP1 (2.7 × 10□ vg); light blue, control uninjected vs. FD uninjected; purple, FD uninjected vs. FD + AAV2.hELP1 (5.4 × 10□ vg); dark blue, FD uninjected vs. FD + AAV2.hELP1 (2.7 × 10□ vg); green, FD uninjected vs. FD + AAV2.eGFP (4.75 × 10□ vg). The control group and FD group shown here is the same as that presented in Figure 1 to allow for direct comparison

### Restoration of functional vision following AAV2.hELP1 gene supplementation

To determine whether the structural and electrophysiological improvements observed with AAV2.hELP1 treatment translated into improvement in visually guided behavior in FD mice, we next assessed visual acuity and contrast sensitivity using the optomotor response (OMR) test. The OMR assay quantifies reflexive head-tracking movements elicited by rotating vertical sinusoidal gratings and provides a noninvasive measure of spatial vision that depends on intact retinal and subcortical visual pathways^33,34^. At 3 months, both AAV2.hELP1 treatment groups (5.4 × 10□ vg and 2.7 × 10□ vg) showed significantly improved visual acuity compared with uninjected FD mice *(p < 0.05; Figure 6Ba)*. By 6 months, only the lower dose group (5.4 × 10□ vg) maintained a significant improvement in visual acuity relative to uninjected FD mice (*p < 0.05; Figure 6Bb*) In terms of contrast sensitivity, FD mice treated with AAV2.hELP1 (5.4 × 10□ vg) demonstrated significantly enhanced performance at 0.061, 0.133, and 0.267 cycles per degree (cpd) in 3 months *(p < 0.05, p < 0.001, p < 0.05, respectively; Figures 6Ca, b, d*). At 6 months, this dose continued to yield significantly greater contrast sensitivity at 0.133, 0.208, and 0.267 cpd compared with uninjected FD mice *(p < 0.05; Figures 6Db-d)*. Together, these improvements in visual acuity and contrast sensitivity align with the structural and electrophysiological rescue observed previously, demonstrating that AAV2-mediated ELP1 gene supplementation effectively restores both retinal and visual pathway function in FD mice.

**Figure 6:**
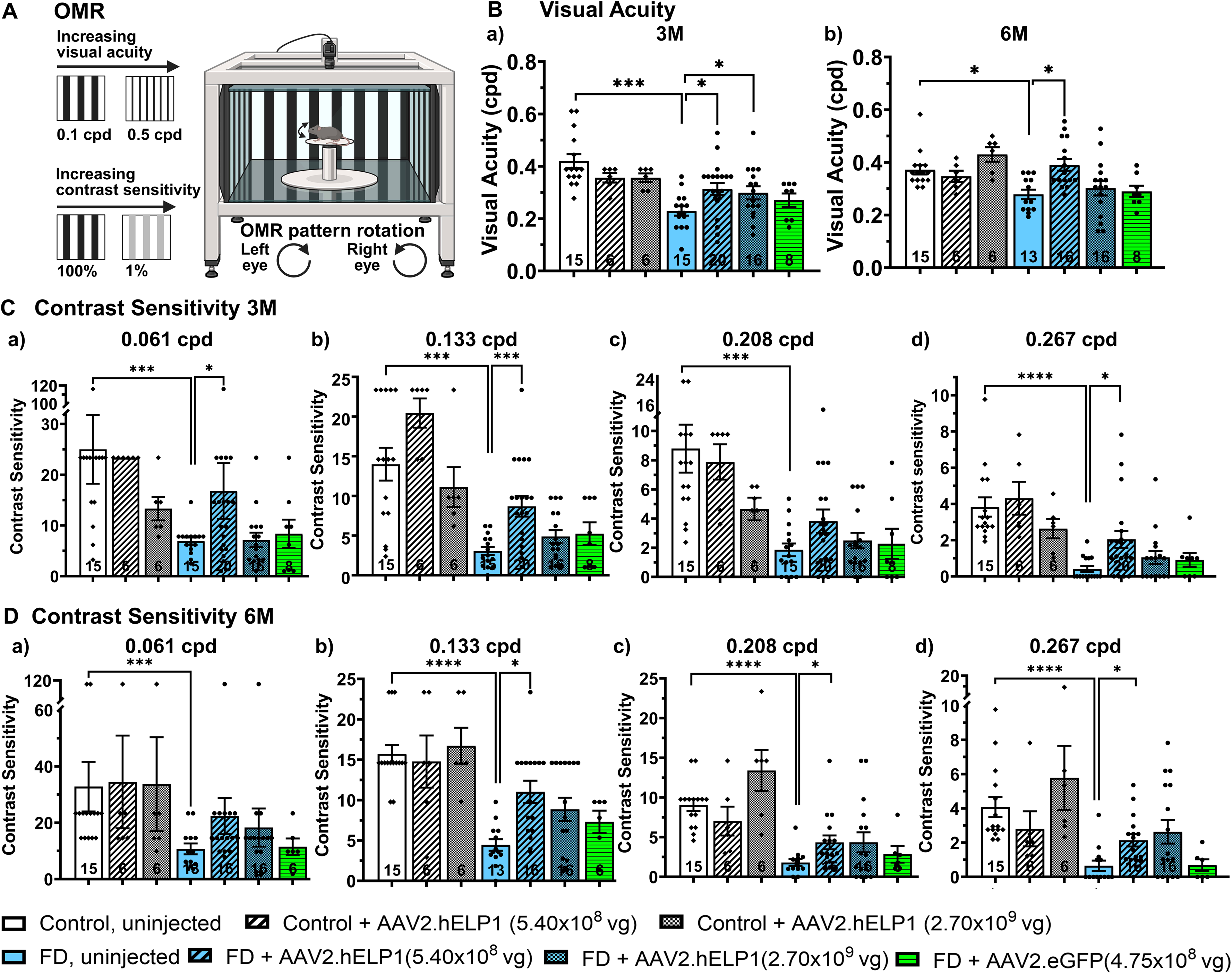
Visual acuity and contrast sensitivity assessed by optomotor response following AAV-mediated gene therapy in FD mice. A. Schematic representation of the optomotor response (OMR) assay used to measure visual acuity and contrast sensitivity. B. Visual acuity measured at (a) 3 and (a) 6 months of age across experimental groups. C. (a-d) Contrast sensitivity measured at 3 months and D. (a-d) 6 months of age at spatial frequencies of 0.061, 0.133, 0.208, and 0.267 cycles per degree (cpd), respectively, indicating age-dependent changes in visual performance following treatment. Data are presented as mean ± SEM and were analyzed using either a t test or Kolmogorov–Smirnov test. Sample sizes for each group are indicated by *n* within each bar; *\*p < 0.05, **p < 0.01, ***p < 0.001, ****p < 0.*0001. The control group and FD group shown here is the same as that presented in Figure 1 to allow for direct comparison

### Intravitreal delivery of AAV2.hELP1 rescues RGC loss in FD mice

Since retinal ganglion cell (RGC) loss is a hallmark of FD and contributes directly to the associated visual deficits, we next investigated whether AAV2-mediated ELP1 gene supplementation could rescue RGC survival by the end of the study period (*Figure. 7A*). As an initial step, we assessed human ELP1 expression in the FD retina to confirm target engagement. Following intravitreal injection of AAV2.hELP1, human ELP1 expression colocalized with Brn3a-positive cells in retinal whole mounts, confirming efficient and targeted human ELP1 transgene expression in RGCs in this study (*Figure. 7C*). Quantitative analysis of RGC density at 6 months post injection revealed a significant increase in RGC survival in FD mice treated with either AAV2.hELP1 (5.4 × 10□ vg) or AAV2.hELP1 (2.7 × 10□ vg), compared with uninjected FD mice *(p < 0.01; Figure. 7B)*. Together, these data establish ELP1 as an essential determinant of RGC survival and provide a mechanistic basis for the observed structural, electrophysiological, and behavioral rescue in FD mice.

**Figure 7:**
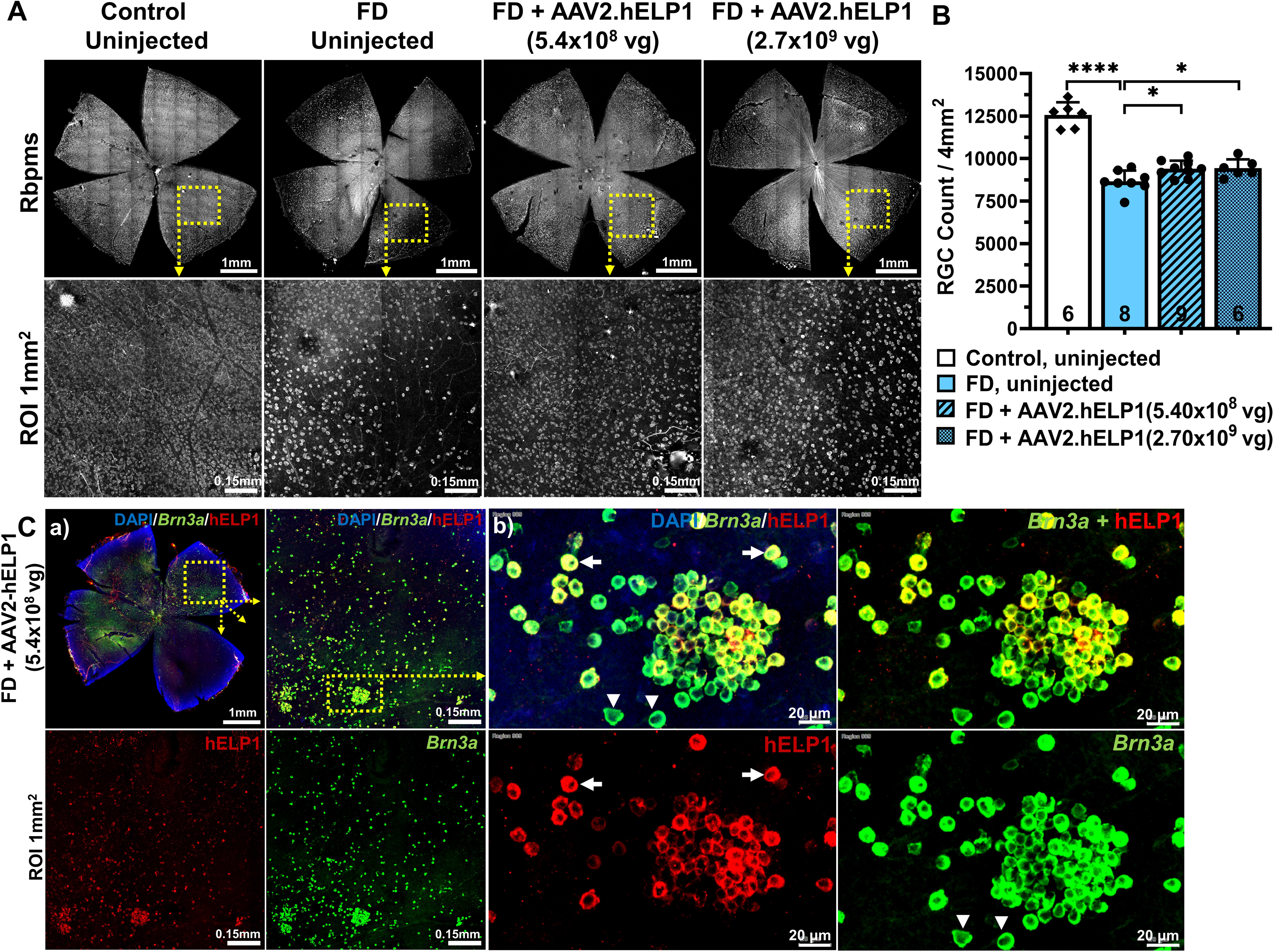
Immunofluorescence analysis and quantification of retinal ganglion cells and localization of human ELP1 in treated retinas. A. Representative retinal flat-mount immunofluorescence images stained for RNA-binding protein with multiple splicing (RBPMS) in P180 control mice and FD mice that were either uninjected or received intravitreal injection (IVI) of AAV2.hELP1 at doses of 5.4 × 10□ vg or 2.7 × 10□ vg. Scale bars, 1 mm (low magnification) and 100 µm (high magnification). B. Quantification of retinal ganglion cell (RGC) density. RGC counts were significantly reduced in FD mice compared with controls and were increased in FD mice treated with either dose of AAV2.hELP1 relative to uninjected FD mice. Cell counts were obtained by enumerating RBPMS-positive cells from four 1 × 1 mm regions per retina. Data are presented as mean ± SEM and were analyzed by one-way ANOVA. Sample sizes for each group are indicated by *n* within each bar; *p* < 0.05, **p** < 0.0001. C. (a,b) Representative immunofluorescence images demonstrating co-localization of Brn3a (green) and human ELP1 (hELP1) in retinal ganglion cells of P30 mice following IVI of AAV2.hELP1 (5.4 × 10□ vg), consistent with expression of the therapeutic transgene in RGCs. Scale bars, 1 mm and 100 µm.

## Discussion

In this study, we present the first comprehensive assessment of retinal structure and function in a retina-specific conditional Elp1 knockout mouse model. Consistent with clinical findings in patients with FD, we observed a significant reduction in RNFL thickness in FD mice compared with control littermates. Interestingly, we also identified swelling of the middle retina and thinning of the outer retina changes (*Figure S1B*). In human retinal disease, typically thickening of the inner nuclear layer (INL), the principal component of the middle retina, is often associated with inflammatory pathology such as diabetic retinopathy or multiple sclerosis related retinopathy and is thought to reflect blood retinal barrier breakdown, microglial activation, Müller cell dysfunction, or retrograde trans-synaptic degeneration^35,36^ Supporting this idea, Schultz et al. reported reactive astrogliosis and microglial activation in Pax6-Cre⁺;Elp1*^loxp/loxp^* mice and in post-mortem FD patient retinae, ^37^suggesting that inflammation may contribute to the middle-retina swelling we observed.

Although Ueki et al. previously reported no gross photoreceptor morphological abnormalities in this model based on histological analysis,^28^ our OCT data revealed measurable thinning of the outer retina, suggesting subtle structural alterations that may occur secondary to RGC degeneration (*Figure S1C*). Consistent with this possibility, indirect photoreceptor loss following RGC injury has been observed in other Elp1-deficient models driven by Tuba1α-Cre^38^. Interestingly, ffERG recordings showed preserved photopic a-wave amplitudes in FD mice, indicating largely intact photoreceptor function yet consistently reduced scotopic and photopic b-waves, reflecting impaired ON-bipolar cell responses. This ERG pattern is characteristic of selective inner retinal dysfunction and supports a model in which primary RGC deficits lead to secondary disruption of upstream synaptic signaling. The absence of a clear reduction in the light-adapted a-wave likely reflects the rod-dominant murine retina, which limits sensitivity to cone-driven differences^39^. Together, these findings suggest that photoreceptor involvement in FD arises predominantly through secondary degenerative mechanisms and further underscore the importance of developing an RGC-specific conditional FD knockout model to dissect potential cell-autonomous roles of ELP1 in retinal physiology.

In our FD mouse model, early molecular and structural deficits predominantly affect RGCs, consistent with other optic neuropathies in which RGC degeneration leads to secondary disruptions in bipolar- and amacrine-cell signaling^40–42^. Electrophysiological analyses further corroborated these deficits: flash VEP amplitudes were consistently reduced, mirroring impaired visual-pathway signaling reported in FD patients (*Figure 4A*), and pattern ERG and phNR responses confirmed pronounced RGC dysfunction (*Figure 4B–C*). Full-field ERG recordings also showed diminished dark- and light-adapted b-waves, indicating impaired inner retinal signaling, while light-adapted a-waves remained largely preserved, consistent with limited sensitivity to cone-driven changes in the rod-dominant murine retina (*Figure 5B*). Together, these findings support a model of trans-synaptic functional impairment across the inner retina driven by primary RGC deficits.

In addition, we demonstrated that AAV-mediated gene therapy using AAV2.hELP1 can rescue both structural and functional visual deficits in FD mice. FD mice treated with AAV2.hELP1 (5.4 × 10□ vg) demonstrated consistent improvements across nearly all measures including OCT, electrophysiology, visual acuity, and contrast sensitivity throughout the 6-month follow-up period compared with uninjected FD mice. In our study, RNFL measurements showed that the response to hELP1 treatment changes over time. The early decrease in RNFL thickness seen at one month after the high-dose injection may be related to the procedure itself or simply reflect the fact that the treatment has not had enough time to counteract ongoing degeneration (*Figure S1A*). By six months, both doses led to thicker RNFL compared with untreated FD mice, indicating that the hELP1 treatment does provide structural benefit as the disease progresses. Notably, the RNFL thickness was increased at six months compared to the three-month time point, suggesting not only attenuation of further thinning but also evidence of axonal remodeling and improved retinal integrity, rather than only slowing of RNFL thinning. These observations clearly demonstrate that ELP1 supplementation rescues RGC integrity over time, and they highlight the importance of considering both dose and timing when developing a gene therapy approach for FD-related optic neuropathy.

Between the two AAV2.hELP1 doses tested, 5.4 × 10□ vg produced the most consistent therapeutic effect. FD mice receiving this dose demonstrated improvements in RNFL and outer retina thickness, reduced middle retina swelling, enhanced flash VEP, pattern ERG, and phNR amplitudes, and superior visual-behavioral outcomes compared with uninjected FD mice. By contrast, FD mice receiving the higher dose (2.7 × 10□ vg) exhibited transiently increased middle-retina thickness at 3 months (*p < 0.01; Figure S1B*) followed by normalization and significant reduction by 6 months (*p < 0.01; Figure S1B*). In the outer retina, this higher-dose group showed decreased thickness at 3 months *(p < 0.05*), suggesting early retinal stress. However, the 5.4 × 10□ vg group displayed significantly greater outer retina thickness at 6 months (*p < 0.05*) compared with uninjected FD mice, indicative of structural recovery (*Figure S1C*). These findings support a dose-dependent effect, where the lower AAV dose achieves durable rescue while the higher dose may transiently exacerbate retinal stress before partial stabilization. Schultz *et al.* similarly observed dose-dependent retinal stress at concentrations exceeding 5.4 × 10□ vg in their studies, supporting this as an optimal therapeutic window^24^.

The improvements in electrophysiological responses following hELP1 treatment are particularly notable because they suggest recovery of retinal circuit function. Functional enhancement within the inner retina is especially meaningful given that RGCs are expected to be the primary cell type transduced by AAV2.hELP1^43,44^. Thus, the improvement in ffERG responses is likely due to better synaptic transmission rather than differences in vector distribution. The restoration of fVEP, pattern ERG, and phNR amplitudes therefore indicates improved trans-synaptic communication from bipolar cells to RGCs and more effective propagation of visual signals along the visual pathway. Collectively, these findings support a model in which stabilizing RGC health through ELP1 supplementation helps preserve the integrity of the broader retinal network, extending therapeutic benefit beyond the most directly impacted cell population.

Confirming a previous study, AAV2.hELP1 delivery also rescued RGC survival. Retinal whole-mount staining demonstrated human ELP1 expression colocalizing with Brn3a-positive cells, confirming RGC-specific transgene expression (*Figure 7C and Schultz et al., 2023*). Quantitative analysis revealed a significant increase in RGC density in FD mice treated with AAV2.hELP1 (5.4 × 10□ vg) or AAV2.hELP1 (2.7 × 10□ vg) compared with uninjected FD mice at 6 months, indicating that preservation of RNFL was accompanied by direct cellular rescue (*Figure. 7B*). This aligns with the functional recovery observed in VEP, pattern ERG, and phNR recordings, indicating that AAV2-mediated *ELP1* supplementation effectively restores both RGC survival and downstream visual signaling.

Finally, this study provides the first comprehensive evaluation of both structural and functional rescue of RGCs in a progressive optic-neuropathy model of FD. Significant anatomical and electrophysiological deficits were observed in the Pax6-Cre⁺;Elp1*^loxp/loxp^* mouse model recapitulating human FD pathology. Intravitreal delivery of AAV2.hELP1 (5.4 × 10□ vg) achieves a robust and sustained rescue of RGC survival, retinal architecture, and visual function, directly supporting AAV2-mediated ELP1 gene supplementation as a clinically viable therapeutic strategy for optic neuropathy and vision loss in familial dysautonomia.

## LIST OF ABBREVIATIONS

AAV2: adeno-associated virus 2;
cpd: cycles per degree;
ELP1: Elongator acetyltransferase complex subunit 1;
ERG: electroretinogram;
FD: familial dysautonomia;
GCIPL: ganglion cell-inner plexiform layer;
hELP1: human EPL1;
IACUC: Institutional Animal Care and Use Committee;
INL: inner nuclear layer;
IPL: inner plexiform layer;
IS: inner segment;
IVI: Intravitreal injection;
OCT: optical coherence tomography;
OMR: optomotor response;
ONL: outer nuclear layer;
OPL: outer plexiform layer;
OS: outer segment;
phNR: photopic negative response;
RGC: retinal ganglion cell;
RNFL: retinal nerve fiber layer;
VA: visual acuity;
VEP: visual evoked potential;
vg: vector genomes

## Availability of data and materials

Materials and datasets used and/or analyzed during the current study are available from the corresponding author upon reasonable request.

## Authors Contributions

A.C. and H.C.C. conceived and designed the study. H.C.C., Y.A., C.E.K., R.E., S.K., M.C., L.H.V., S.A.S., E.M, F.L and A.C. contributed to the development of methodology and to the writing, review, and revision of the manuscript. H.C.C., Y.A., C.E.K., R.E., S.K., M.C., L.H.V., S.A.S., and A.C. were involved in data acquisition, analysis, interpretation, and statistical evaluation. A.C. obtained funding for this study. All authors read and approved of the final manuscript.

## Supplementary figures

**Figure S1: Optical coherence tomography analysis of retinal layer thickness following AAV-mediated gene therapy.**

Experimental groups included control mice with no injection, FD mice with no injection, FD mice receiving intravitreal injection of AAV2.hELP1 (5.4 × 10□ vg), FD mice receiving intravitreal injection of AAV2.hELP1 (2.7 × 10□ vg), and FD mice receiving intravitreal injection of AAV2.eGFP (4.75 × 10□ vg). A. Retinal nerve fiber layer (RNFL) thickness measured by OCT at 1 month of age. B. Middle retinal layer thickness (MRT) measured by OCT at 1, 3, and 6 months of age. C. Outer retinal layer thickness (ORT) measured by OCT at 1, 3, and 6 months of age. Data are presented as mean ± SEM and were analyzed using either a t test or Kolmogorov–Smirnov test. Sample sizes for each group are indicated by *n* within each bar; *\*p < 0.05, **p < 0.01, ***p < 0.001, ****p < 0.0001*.

**Figure S2: Early electrophysiological assessment of visual pathway function following AAV-mediated gene therapy.**

A. Quantification of flash visual-evoked potential (VEP) P2–N1 amplitudes in control and FD mice at 1 month of age across experimental groups. B. Quantification of pattern electroretinogram (pERG) P1–N2 amplitudes at 1 month of age. C. Quantification of photopic negative response (phNR) amplitudes at 1 month of age across experimental groups. Data are presented as mean ± SEM and were analyzed using either a t test or Kolmogorov–Smirnov test. Sample sizes for each group are indicated by *n* within each bar; *p* < 0.05, p < 0.01, *p* < 0.001, p < 0.0001.

**Figure S3: Immunohistochemical analysis of retinal cell types following AAV2.hELP1 treatment in FD mice.**

Representative immunohistochemistry of retinal cross sections from FD mice receiving intravitreal injection of AAV2.hELP1 at doses of 5.4 × 10□ vg or 2.7 × 10□ vg. From left to right: peanut agglutinin (PNA) labeling of cone photoreceptors, rhodopsin labeling of rod photoreceptors, RNA-binding protein with multiple splicing (RBPMS) labeling of retinal ganglion cells, protein kinase C alpha (PKCα) labeling of bipolar cells, glutamine synthetase (GS) labeling of Müller glial cells, glial fibrillary acidic protein (GFAP) labeling of activated Müller glia, calbindin (CALB1) labeling of horizontal cells, and choline acetyltransferase (ChAT) labeling of amacrine cells. Scale bar, 50 µm.

**Figure S4: Full-field electroretinographic analysis following AAV-mediated gene therapy at 1 month.**

Quantification of full-field electroretinogram (ERG) amplitudes, including dark-adapted a-wave and b-wave amplitudes and light-adapted a-wave and b-wave amplitudes, measured at 1 month of age across experimental groups. Data are presented as mean ± SEM and were analyzed by two-way ANOVA. Sample sizes for each group are indicated by *n* within each bar; *\*p < 0.05, **p < 0.01, ***p < 0.001, ****p < 0.0001*.

. The color of the significance markers denotes the comparison group as follows: red, FD + AAV2.hELP1 (5.4 × 10□ vg) vs. FD uninjected; blue, FD + AAV2.hELP1 (2.7 × 10□ vg) vs. FD uninjected; green, FD + AAV2.eGFP (4.75 × 10□ vg) vs. FD uninjected; black, control + AAV2.hELP1 (5.4 × 10□ vg or 2.7 × 10□ vg) vs. control uninjected.

**Figure 5: Representative flat-mount RGC staining following AAV2.hELP1 or AAV2.eGFP injection**

Representative retinal flat-mount immunofluorescence images of retinal ganglion cells (RGCs) stained with RNA-binding protein with multiple splicing (RBPMS) from control mice injected with AAV2.hELP1 (5.4 × 10□ vg) and FD mice injected with AAV2.eGFP (4.75 × 10□ vg).

