## Supplementary figures and images for "ELP1 Gene Augmentation Restores Visual Function in a Mouse Model of Familial Dysautonomia"

### Supplemental figure 1

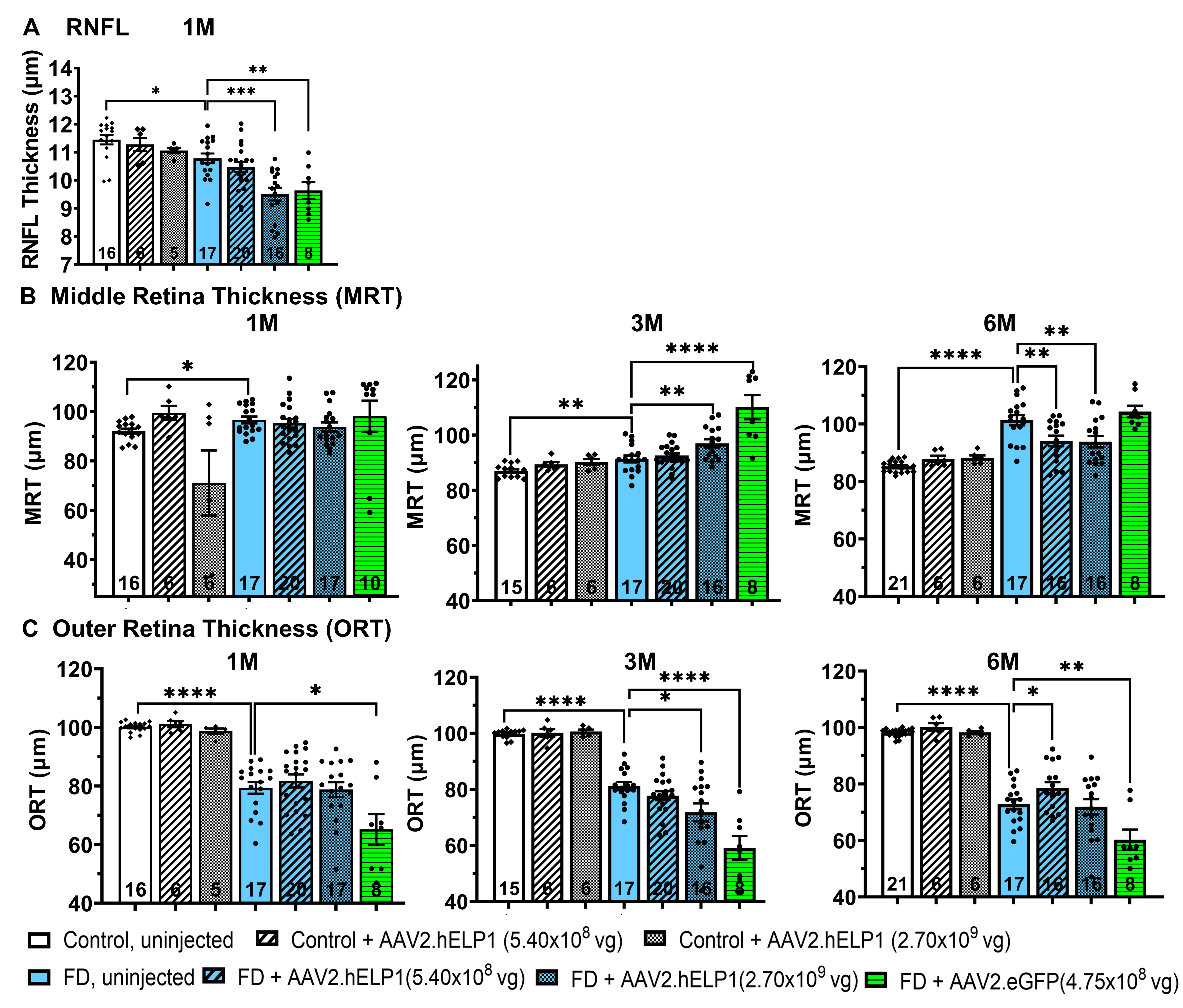

### Supplemental figure 2

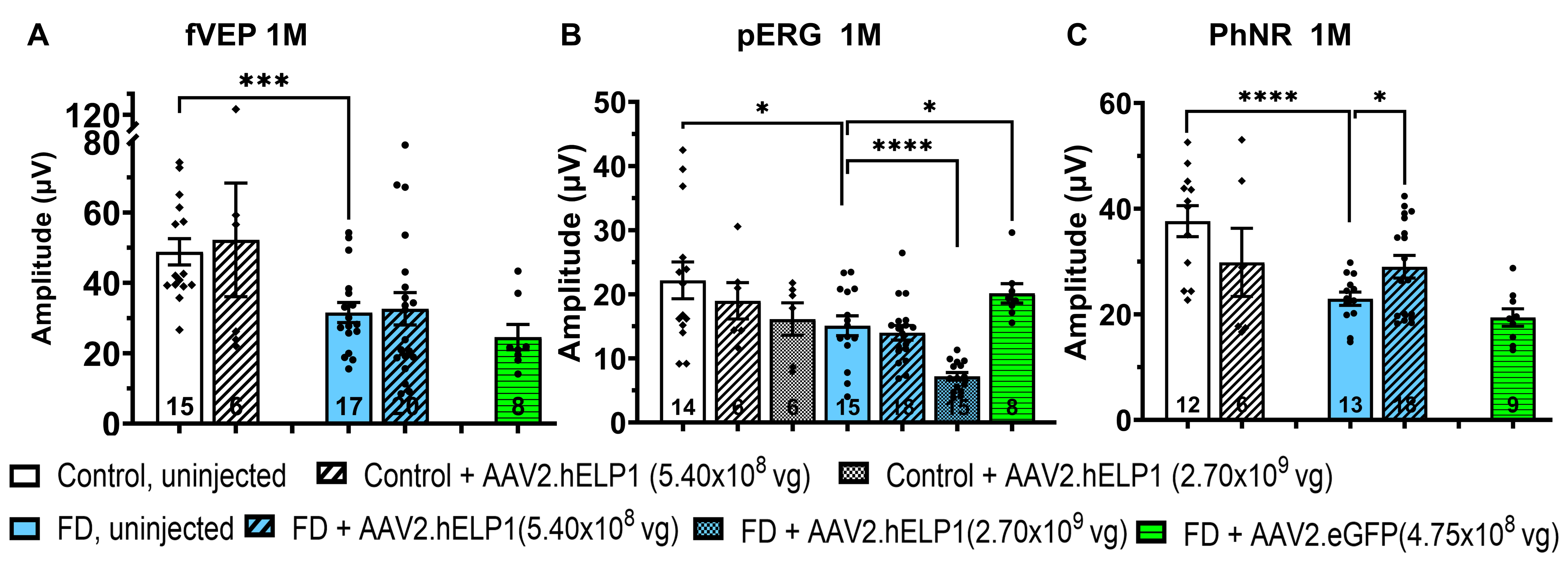

### Supplemental figure 3

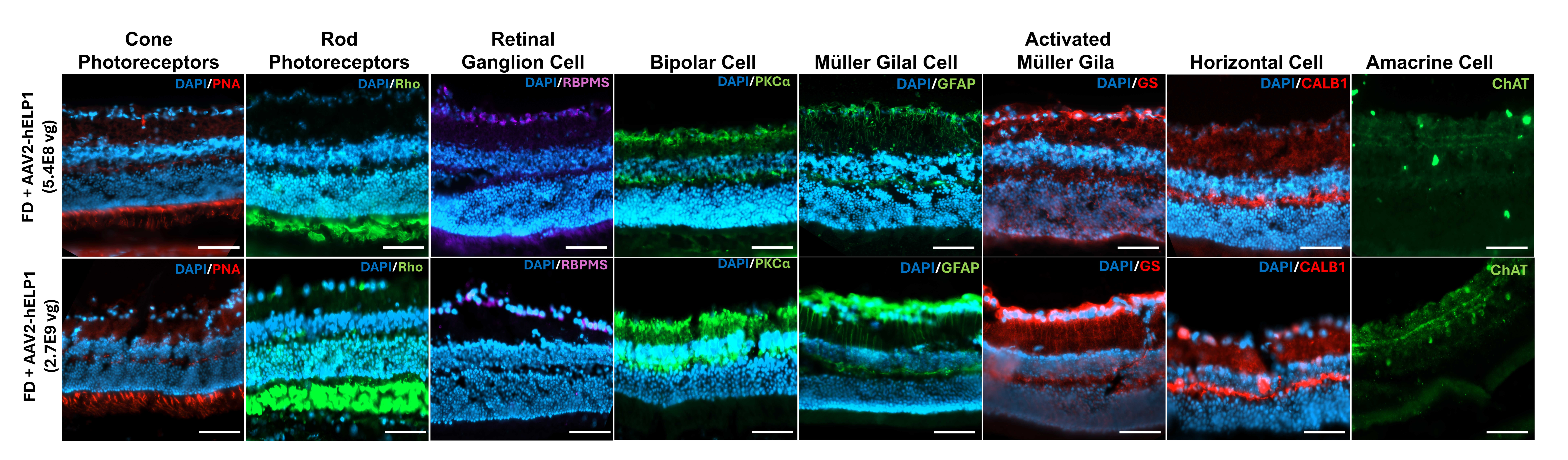

### Supplemental figure 4

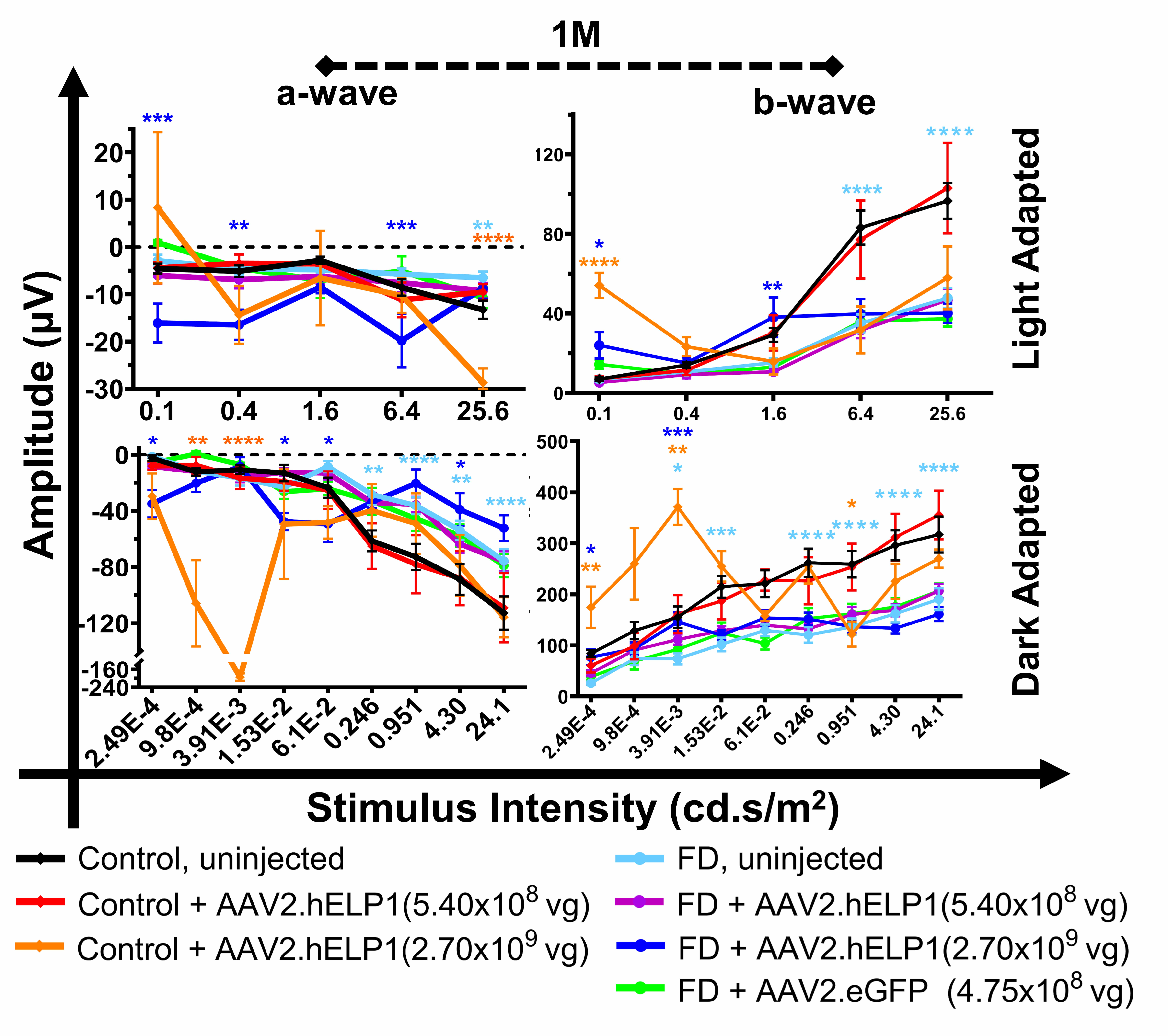

### Supplemental figure 5

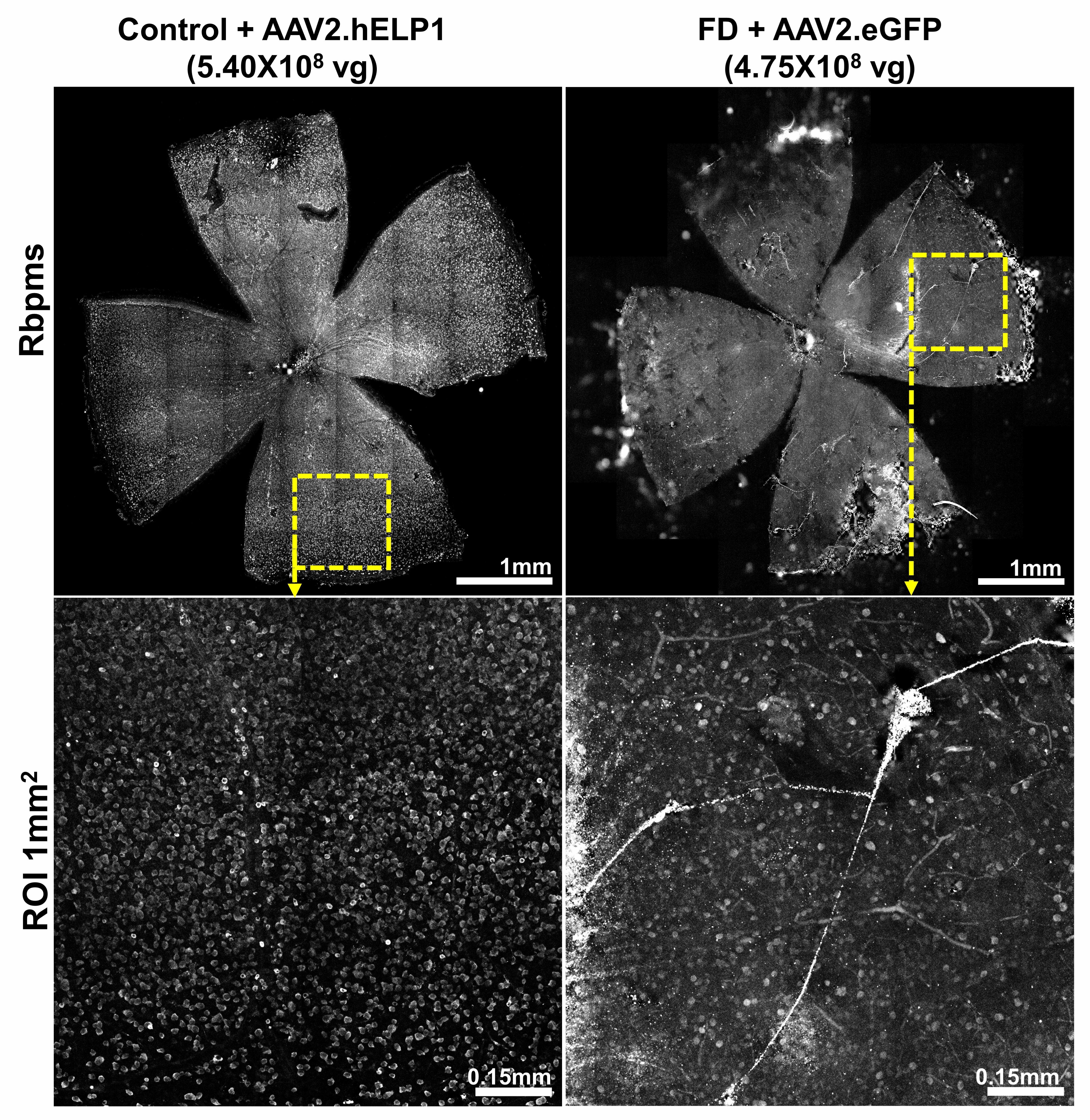
